# Genomic and Dietary Insights from Non-Invasive Sampling of Western Capercaillie in Tyresta National Park (Sweden)

**DOI:** 10.1101/2025.09.24.678326

**Authors:** Tom van der Valk, Julia Hedlund Lindberg, Jonas Ekstrand, Ulf Gyllensten

## Abstract

The Western Capercaillie (*Tetrao urogallus*), a boreal forest specialist, has experienced marked declines in southern Sweden due to habitat loss, fragmentation, and isolation. Tyresta National Park and the surrounding forests harbors one of the region’s last remnant southern populations, yet little is known about its genetic health, population structure, and dietary ecology. Here, we used low-coverage whole-genome sequencing of 273 non-invasively collected samples (feathers and faeces) to comprehensively assess the status of this population. Feather samples yielded high-quality genomic data, enabling sex identification, estimation of genome-wide heterozygosity, runs of homozygosity (ROH), and relatedness inference. Heterozygosity levels in Tyresta were comparable to other European Western Capercaillie populations, and inbreeding levels were generally low, although several individuals showed long ROH indicative of recent close-kin mating. Population structure analyses revealed a genetically distinct and panmictic unit with evidence of female-biased dispersal. Kinship analysis uncovered a highly interconnected social network, with males having on average more relatives than females. Recapture simulations based on genotype matching estimated a current maximum population size of 164 - 208 individuals. Faecal metagenomic analysis revealed a diverse diet, including key plant species such as blueberry, lingonberry and alder, and a wide range of invertebrates critical for chick development. Despite its current genetic robustness, the Tyresta population is small, isolated, and at risk of inbreeding depression. Conservation efforts should prioritize maintaining habitat quality, promoting gene flow, e.g. by protecting migration corridors to neighbouring populations and ensuring a rich forest invertebrate community to support population resilience.

## 1. Introduction

The western capercaillie (*Tetrao urogallus*) is the largest grouse species in Europe and a flagship species of boreal and montane coniferous forests across Eurasia (Artfakta 2020). In Sweden, it is primarily associated with old-growth pine (*Pinus sylvestris*) and spruce (*Picea abies*) forests, where it relies on a structurally complex habitat mosaic that includes dense cover for roosting, open ground for lekking, and a diverse understory of blueberry (*Vaccinium myrtillus*) for foraging (Angelstam 2004). Despite its cultural and ecological significance, capercaillie populations have undergone dramatic declines across much of their range due to habitat fragmentation, forest management practices, climate change, and increased human disturbance (“European Red List of Birds” 2021). These pressures have led to reduced population connectivity, increased isolation, and heightened conservation concern, particularly in fragmented landscapes such as southern Sweden (Schimmel and Granstrom 1996).

Tyresta National Park, located just outside Stockholm, represents one of the few remaining patches of old-growth boreal forest in the southern part of the country. Parts of the forest have never been logged and trees up to 350 years old have been recorded. Despite its proximity to urban areas, Tyresta supports a small, isolated population of capercaillies, whose long-term viability is uncertain. The Tyresta National Park and the neighboring Tyresta Nature Reserve together encompass around 5,000 hectares of forested land. A devastating wildfire in 1999 affected 450 hectares of the central area, and created a heterogeneous post-fire landscape (Schimmel and Granstrom 1996). While such disturbances can create early successional habitats beneficial to some species, they may also impact capercaillie habitat quality and availability. In the case of the 1999 fire, it destroyed key habitats and lekking areas, and likely substantially reduced the population. Understanding the status and genetic health of the capercaillie population in Tyresta is crucial for informing local conservation strategies and for contributing to broader efforts aimed at reversing declines in southern Sweden.

Effective management and conservation of the capercaillie depend on accurate assessments of population size, connectivity, and genetic diversity. However, traditional monitoring approaches, such as capturing and marking individuals, are logistically challenging, costly, and potentially harmful to this elusive and sensitive species (Rolf Brittas, Endre Hofstad Hansen, Torfinn Jahren, Marius Kjønsberg, Mikkel Andreas Jørnsøn Kvasnes, Eric Ringaby, Maria Hörnell-Willebrand August-2024). Non-invasive genetic sampling, using shed feathers and faeces, provides a powerful alternative that minimizes disturbance while enabling broad-scale and repeatable sampling across time and space (Taberlet, Waits, and Luikart 1999; Carroll et al. 2018). However, such samples often yield low quantities of degraded DNA and are contaminated with high levels of environmental and microbial DNA, complicating genomic analysis.

Advances in high-throughput sequencing and bioinformatics have dramatically improved our ability to recover genome-wide data even from low-quality non-invasive samples. In particular, low-coverage whole-genome sequencing (lcWGS) has emerged as a cost-effective approach for studying wildlife populations at the genomic level (Perry et al. 2010). By sequencing many individuals at shallow depth, lcWGS enables estimation of key metrics such as genome-wide diversity, inbreeding, population structure, and kinship, parameters that are essential for assessing population resilience, identifying signs of inbreeding depression or genetic drift, and informing targeted conservation actions (Shafer et al. 2015; van der Valk and Dalèn 2024).

In this study, we applied lcWGS to a set of 273 non-invasively collected capercaillie samples from Tyresta National Park and the surroundings (**Figure 1**). Our objectives were to: (1) evaluate the efficacy of different sample types (feathers and faeces) for generating genomic data; (2) assess the genetic health of the population by quantifying levels of genome-wide heterozygosity and inbreeding; (3) investigate population structure and connectivity within the park; and (4) reconstruct kinship relationships to infer patterns of relatedness and potential family structure within the population. Additionally, we explored the metagenomic content of faecal samples to infer dietary composition, providing further ecological context to support holistic conservation efforts (Soininen et al. 2009).

**Figure 1.**
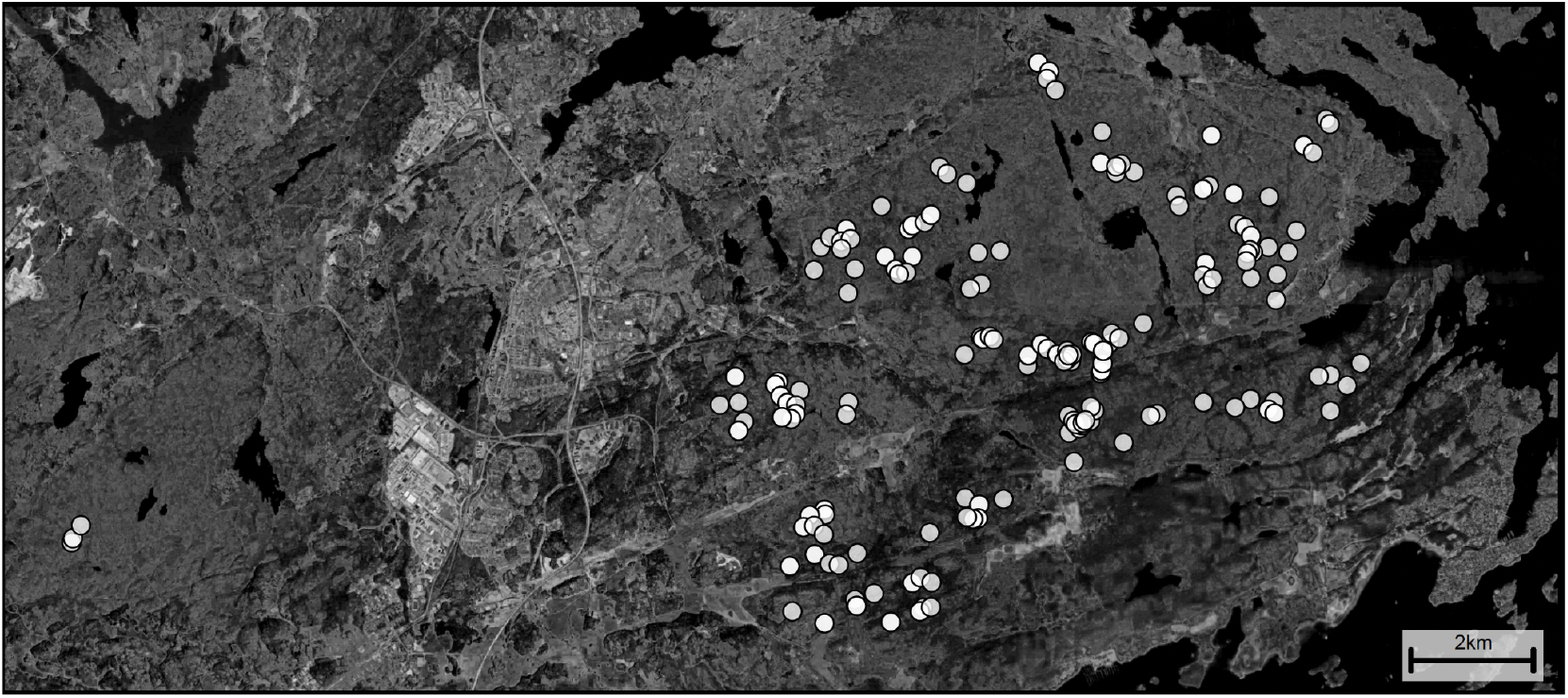
Geographic location of the sequenced samples. Map of Tyresta National Park showing the collection sites of capercaillie samples. Map is displayed in grey-scale to obscure the exact locations of the leks.

## 2. Methods

### 2.1. Sample Collection and Sequencing

A total of 273 samples were collected non-invasively during April-June 2022 and March-September 2023 (Supplementary Table S1), consisting of shed feathers and faecal pellets. Samples were handled with gloves and collected in individual plastic bags labeled with GPS coordinates of their respective location. The material was temporarily kept at −20°C until shipped on dry ice to Uppsala University for registration with serial number barcodes and final storage at −80°C.

DNA extractions were performed in a lab dedicated to pre-PCR work where each feather or fecal pellet was handled individually, using clean forceps and sterile scalpels on a cutting mat cleaned with ethanol (70%) in between samples. Smaller size feathers were used as a whole if they would fit in a 2 mL tube, and from larger feathers approximately 1.5 cm of the basal end were used and cut into smaller pieces to facilitate lysis. The largest wing- or tail feathers were divided longitudinally for dissection of the superior umbilicus, aiming to reach remnants of blood cells of the axial artery as described by Horváth et al (2005), and when possible, the basal tip of the calamus were also included. DNA extraction from feathers was performed using the DNeasy Blood & Tissue Kit (Qiagen, cat.no. 69504) following a user-developed protocol with minor modifications (https://www.qiagen.com/us/resources/resourcedetail?id=a5a065dc-e287-4a61-b917-9792e25ab42f&lang=en). Feathers were transferred to 2 mL DNA LoBind microcentrifuge tubes (VWR, cat.no. 525-0131) containing 300 µL ATL buffer, 20 µL proteinase K and 20 µL 1M DTT (Merck, cat.no. 43816-10ML) and vortexed for 10 seconds before an overnight incubation at 56°C with constant shaking at 550 rpm on a Thermomixer (Thermo Fisher Scientific Inc.). The lysed samples were centrifuged at max speed for 30 seconds and transferred to new 2 mL tubes avoiding pipetting any remaining solid material. A mix of 300 µL AL buffer and 300 µL ethanol (100%) was added, and samples thoroughly vortexed followed by a short spin. Lysates were transferred to labeled DNeasy Mini spin columns in smaller aliquotes of maximum 600 µL and centrifuged at 6 000 x *g* for 1 minute on a benchtop Eppendorf® centrifuge with discarding of the flow-through. When the total volume of lysate had passed through the columns, membranes were washed with 500 µL AW1 buffer and centrifuging 1 minute at 6 000 x g and discarding of the flow-through, followed by a final wash with 500 µL AW2 buffer, centrifuging at 20 000 x *g* for 6 minutes and discarding of the flow-through. The columns were moved to new, labeled 1.5 mL DNA LoBind Eppendorf tubes and 50 µL AE buffer pre-warmed to 70°C were added to the membranes to incubate at room temperature for 5 minutes. Finally, total DNA was eluted by centrifugation at 6 000 x *g* for 1 minute. To increase DNA recovery, the elution was performed twice resulting in two separate 50 µL aliquots from each sample. DNA from the faecal samples were extracted using QIAamp Fast DNA Stool Mini Kit (Qiagen cat.no. 51604) according to the manufacturers protocol *Isolation of DNA from Stool for Human DNA Analysis* with some modification described by Vallant et al. (2018). Approximately 100 mg of the outer parts of the frozen dropping were cut into smaller pieces avoiding visible traces of uric acid. The faecal samples were transferred to 2 ml DNA LoBind Eppendorf tubes containing 1 mL InhibitEX buffer pre-warmed to 70°C the samples were homogenized by slightly grinding using the end of a sterile and DNA-free swab (Sarstedt cat.no. 80.626). Samples were vortexed for 1 minute and incubated on a Thermomixer at 56°C for 1 hour with shaking at 550 rpm, before centrifugation at max speed for 6 minutes. A volume of 600 µL of each sample were pipetted into new DNA LoBind tubes containing 25 µL proteinase K, and 600 µL AL buffer were added and samples vortexed for 15 seconds before incubation on a Thermomixer at 56°C over-night under constant shaking at 550 rpm. The lysed material was spun briefly before adding 600 µL Ethanol (100%) and 4 ng carrier RNA and mixed by vortexing. Isolation of DNA was performed by adding 600 µL aliquotes of lysate to labeled QIAamp spin columns and centrifuging at maximum speed for 1 minute and discarding the flow-through. When the total volume of lysate had passed through the columns, membranes were washed with 500 µL AW1 and centrifuging at maximum speed for 1 minute with discarding of the filtrate, followed by a second wash with 500 µL AW2 buffer, centrifuging at 10 000 x g for 5 minutes and discarding of the filtrate. Lastly, the spin columns were moved to new 2 ml tubes and centrifuged 1 min at maximum speed before being transferred to new, labeled 1.5 µL DNA LoBind Eppendorf tubes and 50 µL ATE buffer pre-warmed to 70°C. The samples were incubated at room temperature for 5 minutes followed by centrifugation at 8 000 x g for 1 minute to elute the DNA. To increase DNA recovery, the elution was performed twice resulting in two separate 50 µL aliquots from each sample. DNA concentrations and quality were measured with Qubit 4 fluorometer (Thermo Fisher Scientific Inc.) using the Qubit 1X dsDNA HS Assay Kit (Thermo Fisher Scientific, cat.no. Q33231), and the Nanodrop 1000 spectrophotometer (Thermo Fisher Scientific Inc.). Sequencing libraries were prepared from 0,001 – 100 ng DNA at the SNP&SEQ Technology Platform in Uppsala, using the in-house developed SPLAT protocol with exclusion of the sodium bisulfite conversion of DNA (Raine et al. 2017). Sequencing was performed using Paired-end 150bp read length, on the NovaSeq X Plus system, using the 10B flow cell and XLEAP-SBS sequencing chemistry, at the SciLifeLab National Genomics Platform in Uppsala. Additionally, we included sequencing data of previously published capercaillie genomes (ENA: PRJEB58761) from Austria (n=5), Bulgaria (n=5), Slovakia (n=4) and northern Sweden (n=1) as a comparative outgroup set.

### 2.2. Data Processing and Quality Assessment

Raw sequencing reads were processed to remove sequencing adapters and low-quality bases using AdapterRemoval v2.3.4 (Schubert, Lindgreen, and Orlando 2016). The cleaned reads were then mapped to the capercaillie reference genome (bTetUro1.1) with bowtie2 v.2.5.4 on the –very-sensitive settings (Langmead and Salzberg 2012) and reads below a length of 40 basepairs or a mapping quality of 20 were discarded. PCR duplicates were filtered out with samtools v.20 markdup (Li et al. 2009). We then assessed sample quality by calculating the percentage of endogenous DNA (reads mapping to the capercaillie genome) and the mean sequencing depth across the genome using samtools v1.20 idxstats and samtools v1.20 depth. Out of the 273 samples, 203 were sequenced above 0.25X coverage, which were then further analysed.

### 2.3. Sex Determination

In birds, females are the heterogametic sex (ZW) and males are homogametic (ZZ). We determined the sex of each individual by calculating the ratio of the average sequencing depth on the Z chromosome to the average depth on the autosomes using samtools v1.20 depth. Males (ZZ) are expected to have a ratio of∼1, while females (ZW) are expected to have a ratio of∼0.5, as they have twice as many autosomes as Z chromosomes.

### 2.4. SNP calling

We called single nucleotide polymorphism (SNPs) across all genomes above 0.25X in coverage using bcftools v1.20 on default parameters (Danecek et al. 2021). We then filtered out sites with more than two alleles, indels, SNPs with more than 50% missing data and those with a minimal allele count below 5.

### 2.5. Genetic Diversity and Inbreeding

Genome-wide heterozygosity, the proportion of heterozygous sites in an individual’s genome, was calculated for each sample with sufficient data coverage (>5X). We used samtools v1.20 mpileup to sample all mapped bases at each genomic site and calculated the fraction of sites above depth 5 for which the minor base frequency at that site was at least 5%. To assess recent inbreeding, we identified Runs of Homozygosity (ROH), which are contiguous stretches of homozygous genotypes in the genome. To call ROH across the complete genome we first imputed missing genotypes using Beagle v5.5 on default settings for all genomes above 0.25X (Browning et al. 2021). We then identified ROH using bcftools v1.20 roh on default settings (Xue et al. 2015) and filtered out ROH with a phred-score below 30.

### 2.5. Population Structure Analysis

To investigate genetic relationships and population subdivision, we performed a Principal Component Analysis (PCA) on the genomic data for all samples above 0.25X coverage using emu v1.2.1 (Meisner et al. 2021). We also constructed a haplotype network based on the near complete mitochondrial DNA (mtDNA) sequences, to visualize relationships among individuals and maternal lineages, respectively. Mitochondrial consensus sequences were obtained using ANGSD v0.940 -dofasta, selecting the major base at each covered mtDNA site (Korneliussen, Albrechtsen, and Nielsen 2014). The haplotype network was constructed using PopArt (Leigh and Bryant 2015)

### 2.6. Genotype Imputation and Kinship Analysis

For kinship analysis we used READ v2.1.1, which can accurately assign up to 3rd degree relationships on low-coverage genomes (Alaçamlı et al. 2024). First, random alleles were called at each genomic position for each individual above 0.25X in coverage using ANGSD v0.940 (-uniqueOnly 1 - remove_bads 1 -minMapQ 30 -minQ 30 -doCounts 1 -dohaplocall 1 -minMinor 1). These allele calls were then converted to plink’s BED/BIM/FAM format and we subsequently used READ v2.1.1 on default parameters to identify related individuals, including identical individuals, parent-offspring pairs, full siblings, and second or third-degree relatives.

### 2.7. Population size estimates

To estimate the population size of capercaillie in Tyresta National Park and its surroundings, we simulated populations of varying sizes, ranging from 10 to 400 individuals. From each simulated population, we randomly sampled 203 birds with replacement (i.e. the number of genomes sequenced above 0.25X), repeating this process 10,000 times. For each iteration, we recorded the frequency with which individual birds were resampled and compared the resulting distribution across the 10,000 simulations to our empirical observations. All simulations were conducted in Python, and the code is available at: https://github.com/tvandervalk/capercaillie_simulations.git

### 2.8. Dietary Analysis

For the faecal samples, reads that did not map to the capercaillie genome were used for metagenomic analysis. To identify plant, fungal, and (in)vertebrate DNA present in the faeces, providing insights into the birds’ diet and gut microbiome, non-host reads were taxonomically classified using kraken2 v2.14 and the full NT database v20240530 (Wood, Lu, and Langmead 2019). Species with the highest number of unique assigned kmers across all faecal samples were then identified using a custom python script. To obtain seasonal insights into plant consumption, all non-host reads were additionally aligned to a database of 15,235 complete chloroplast genomes using Bowtie2, retaining the best match for each read. To reduce redundancy, only a single chloroplast genome per genus was kept, selecting the genome with the highest read support (as reads often aligned to multiple species within a genus). Finally, faecal samples were grouped by collection date to assess temporal changes in plant dietary composition.

## 3. Results

### 3.1. Sequencing Yield and Sample Quality

From the 273 non-invasively collected samples, we successfully generated low-coverage whole-genome sequencing data above 0.25X for 203 samples (**Figure 1**). The sequencing quality and endogenous DNA content varied significantly between sample types. Feather samples generally yielded higher autosomal coverage compared to faecal samples, with many feather samples achieving over 5X coverage and some exceeding 10X, while most faecal samples remained below 5x coverage (**Figure 2A**). Despite the low and variable coverage, we could confidently determine the sex of the individuals. By calculating the ratio of sequencing depth on the Z chromosome to that on the autosomes, we clearly distinguished between males (ratio∼1.0) and females (ratio∼0.5) (**Figure 2B**). This method proved effective even for samples with very low endogenous DNA content (**Figure 2B**). One individual was identified with a potential chromosome aneuploidy, showing a ratio of∼2.0 (**Figure 2B**).

**Figure 2.**
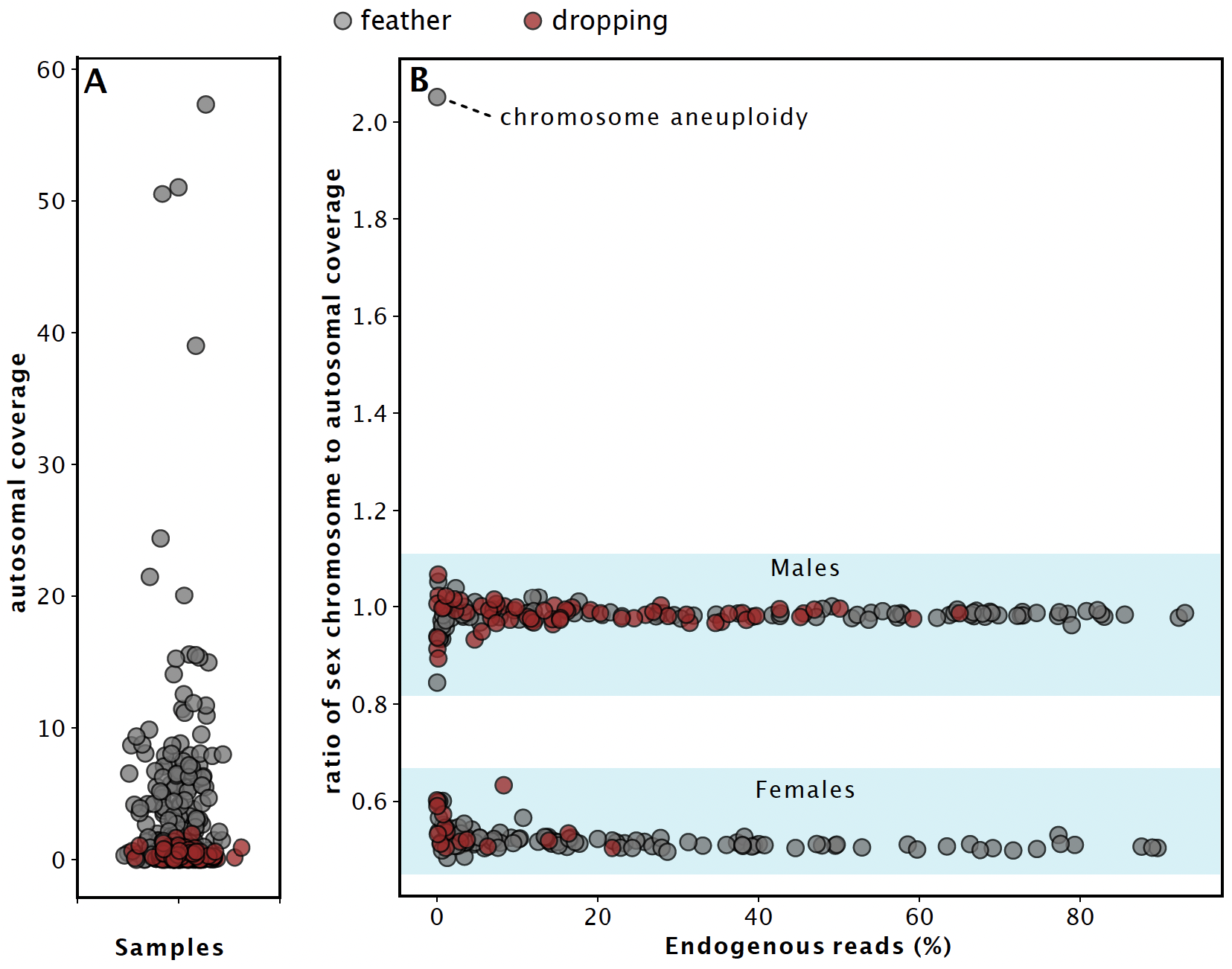
Sequencing Quality and Sex Determination from Non-invasive Samples. This figure illustrates the quality of genomic data obtained and the method for sex determination. **(A)** Plot of the mean autosomal sequencing coverage achieved for each of the 273 samples, categorized by source type. Feather samples (grey) consistently yielded higher coverage than faecal droppings (red). **(B)** Scatter plot showing sex identification based on the ratio of Z chromosome coverage to autosomal coverage versus the percentage of endogenous DNA. Males (ZZ) cluster around a ratio of 1.0, while females (ZW) cluster around 0.5, allowing for accurate sexing even at very low concentrations of endogenous DNA. One individual showing a signal of chromosomal aneuploidy is noted.

### 3.2. Genetic Diversity and Inbreeding

We assessed the genetic health of the Tyresta capercaillie population by analyzing genome-wide heterozygosity and runs of homozygosity (ROH). For individuals with sufficient genomic data (over 1 million sites with a depth >5), autosomal heterozygosity ranged from approximately 0.0025 to 0.0095 (**Figure 3A**). The Tyresta population’s heterozygosity levels were comparable to those observed in outgroup samples from other European populations (**Figure 3A**). Inbreeding levels were evaluated by quantifying the total length of the genome found in ROH. Overall, the Tyresta population exhibited low levels of inbreeding with the distribution of ROH similar to that of outgroup populations from Austria, Bulgaria, and Slovakia (**Figure 3B**). However, a few individuals, both males and female, within the Tyresta population showed notable signs of recent inbreeding, with a significant portion of their genome composed of long ROH segments (>1Mb), indicating recent parental relatedness (**Figure 3B, Supplemental Figure 1**).

**Figure 3.**
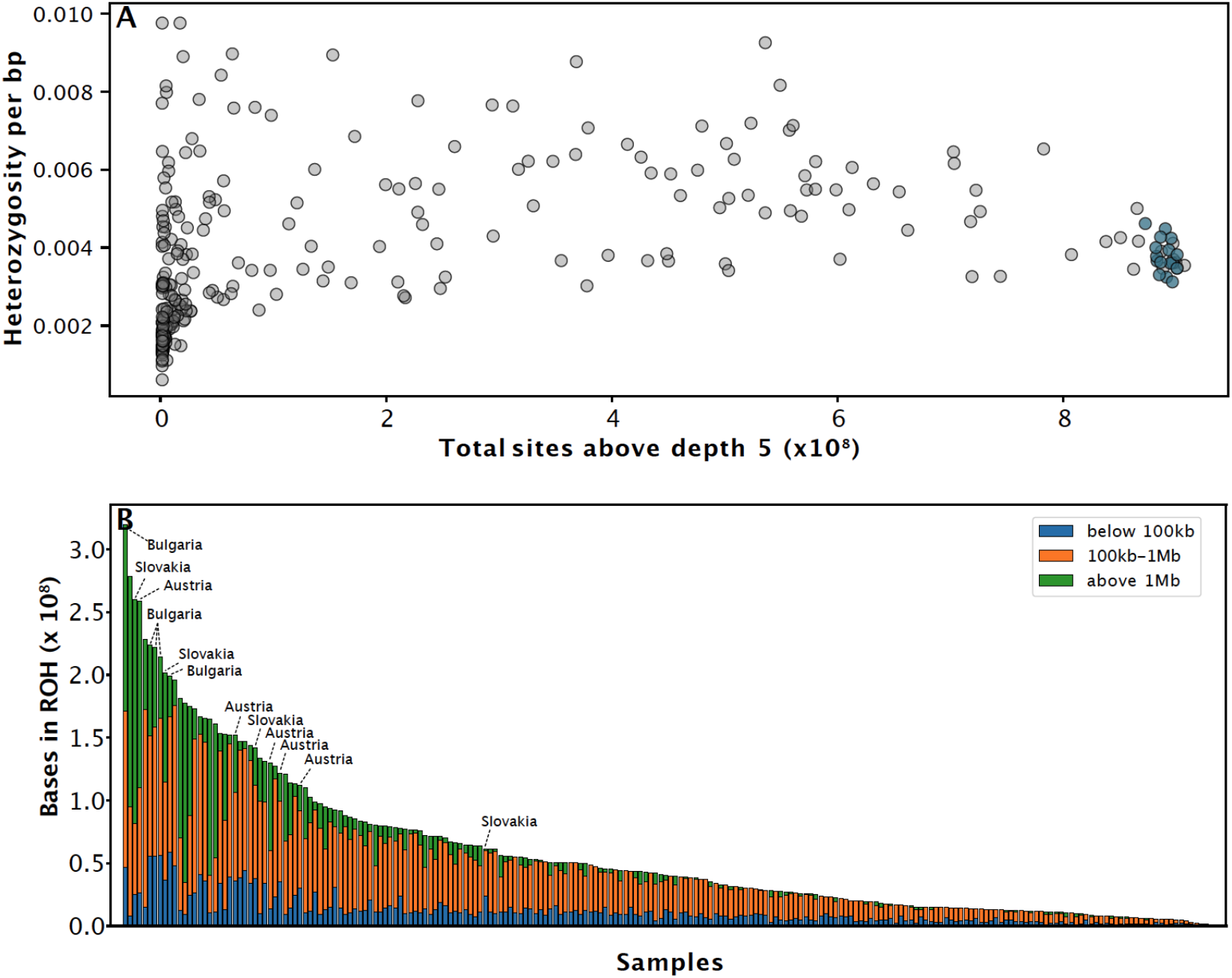
Genetic Diversity and Inbreeding in the Tyresta Capercaillie Population. **(A)** Autosomal heterozygosity per base pair against the total number of genomic sites sequenced at a depth greater than 5X. Samples from Tyresta (grey) show heterozygosity ranging between 0.0025 and 0.0095 for well-covered samples, while outgroup samples are shown in light blue. **(B)** Stacked bar chart quantifying inbreeding through the total length of the genome found in Runs of Homozygosity (ROH). Each bar represents a single bird, with segments color-coded by the length of homozygous blocks: short (<100kb, blue), medium (100kb–1Mb, orange), and long (>1Mb, green). Overall inbreeding in the Tyresta population is low, though some outgroup individuals show significantly higher levels of inbreeding

### 3.3. Population Structure and Kinship

To investigate the genetic structure within Tyresta National Park, we performed a principal component analysis (PCA) on the genomic data (**Figure 4A**). The PCA revealed that the Tyresta capercaillies mostly form a single, relatively homogeneous genetic cluster. The Tyresta cluster is distinct from the outgroup populations from Bulgaria, Slovakia/Austria, and another Swedish population from Jämtland (**Figure 4A**). No clear sub-structure was observed within the Tyresta population itself, although several individuals, of which 75% are male, appear to be slight outliers, potentially representing recent immigrants (**Figure 4A**). Analysis of mitochondrial DNA haplotypes further supported this finding, showing a connected network without deeply divergent maternal lineages, which is consistent with a single, cohesive population (**Figure 4B**). Next, kinship analysis was performed to reconstruct a pedigree and understand relatedness patterns within the population. The analysis identified numerous relationships, including identical individuals (i.e., recaptures), parent-offspring pairs (1st degree), full siblings (2nd degree), and more distant relatives (3rd degree) (**Figure 5**). A network analysis showed extensive connections among individuals up to the 3rd-degree level, confirming the population is highly interconnected (**Figure 5**). Except for 11 individuals, all other 192 samples had at least one other (up to third degree) relative among the samples set. The average individual had∼8 up to third degree relatives in the sample set, with several of the individuals having over 20 relatives. When comparing the number of relatives by sex, males were found to have, on average, a higher number of relatives within the park than females (8.5 against 7.6 respectively), albeit not significant (p∼ 0.11) (**Figure 6**). To assess whether relatedness is reflected in spatial patterns, we compared the Euclidean distances between geographic sampling locations by sex and degree of relatedness (**Figure 7**). As expected, identical individuals (recaptures) were located significantly closer to each other than any other category. Most identical male samples were found within 1 km of each other, suggesting this distance may approximate the territory size of males, although some outliers were detected up to 5 km apart. Among related individuals, females tended to occur in closer proximity than males, consistent with sex-biased differences in dispersal. On average, male relatives were found farther apart than female relatives, suggesting that male capercaillies may disperse over larger spatial scales within the park (**Figure 7**).

**Figure 4.**
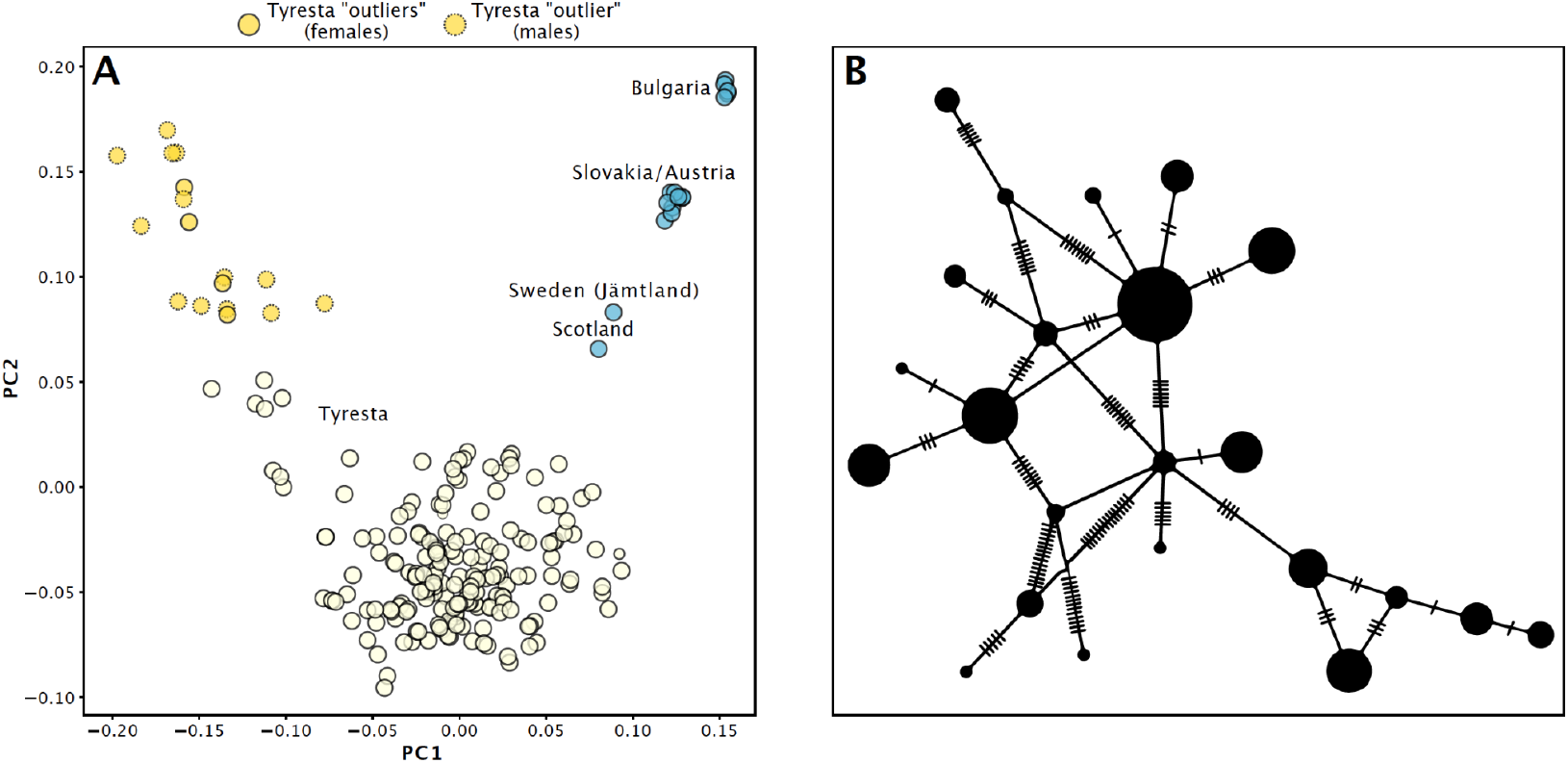
Population structure analysis of capercaillie. **(A)** Principal Component Analysis of genomic data from Tyresta individuals (yellow-green) and outgroup populations (light blue). The Tyresta samples mostly form a single, dense cluster, indicating a lack of substructure, with several individuals in the top left possibly being immigrants. **(B)** Mitochondrial DNA haplotype network. Each circle represents a unique haplotype, with its size proportional to its frequency. The hashes on the lines between haplotypes represent mutational steps. The network shows multiple, interconnected haplotypes, indicating widespread maternal lineages and a lack of population substructure.

**Figure 5.**
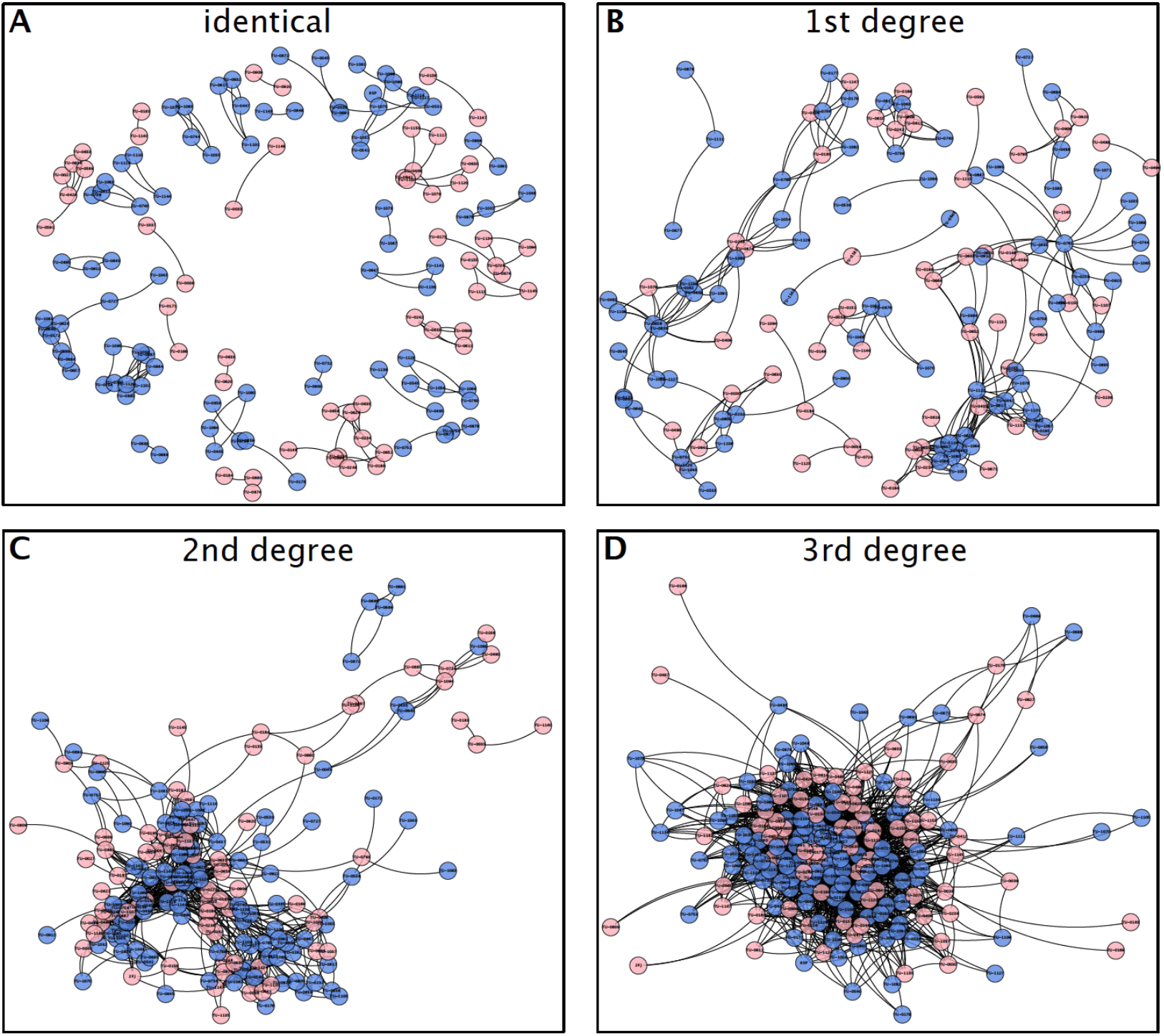
Network Visualization of Kinship Within the Tyresta Population. This figure displays the social structure of the capercaillie population through network graphs, with each panel illustrating a different degree of relatedness. Nodes represent individual birds (males in blue, females in pink). **(A)** Identical: Links connect samples originating from the same individual. **(B)** 1st Degree: Shows parent-offspring and full-sibling relationships. **(C)** 2nd Degree: Shows relationships such as half-siblings or grandparent-grandchild. **(D)** 3rd Degree: Shows connections between cousins and other more distant relatives.

**Figure 6.**
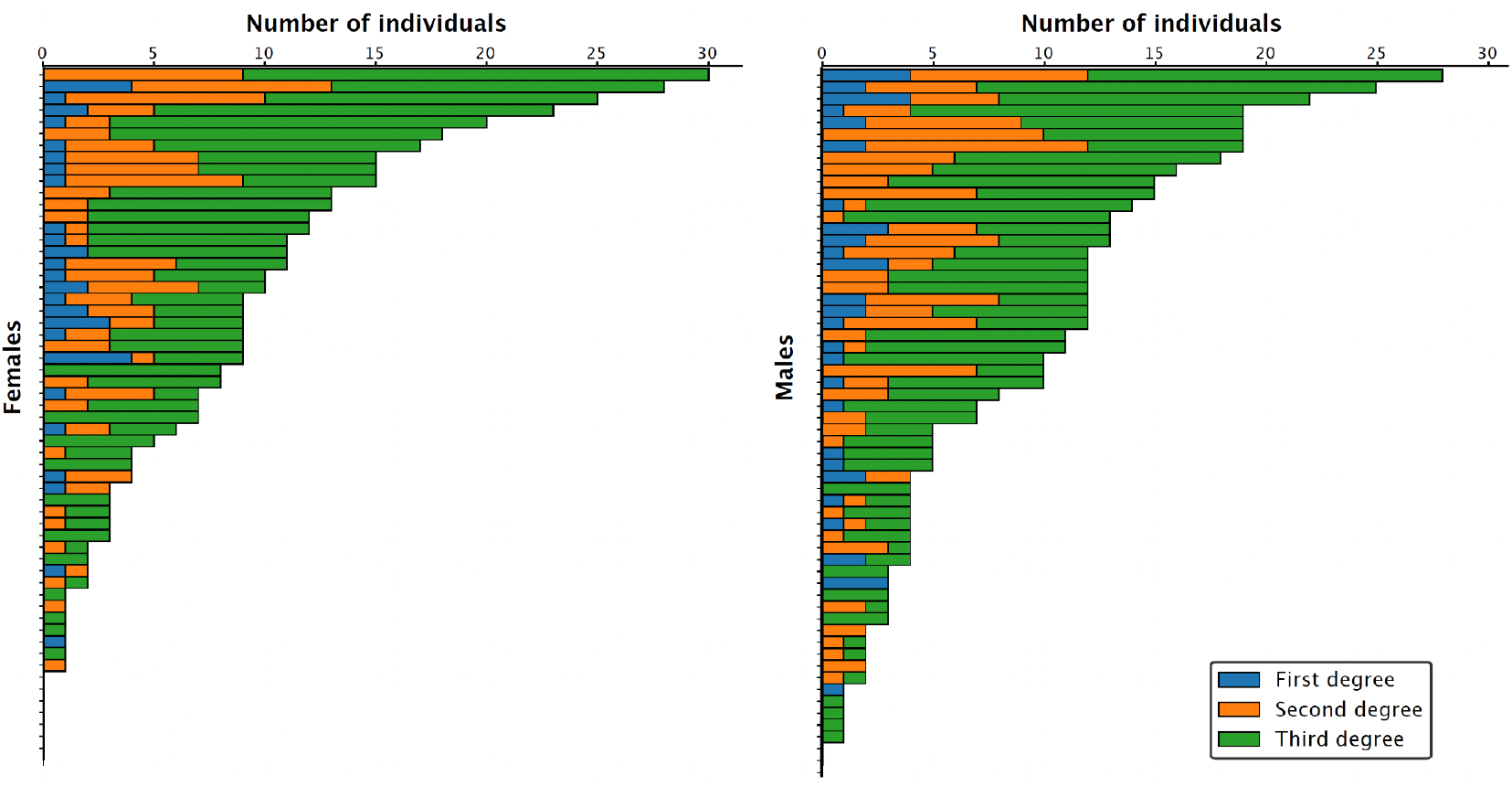
Kinship Breakdown for Each Individual. This chart details the number and degree of relatives for each individual bird sampled. Each horizontal bar represents a unique individual. The color-coded segments quantify the number of relatives of a specific degree that were identified in the study population: first-degree (blue), second-degree (orange), and third-degree (green).

**Figure 7.**
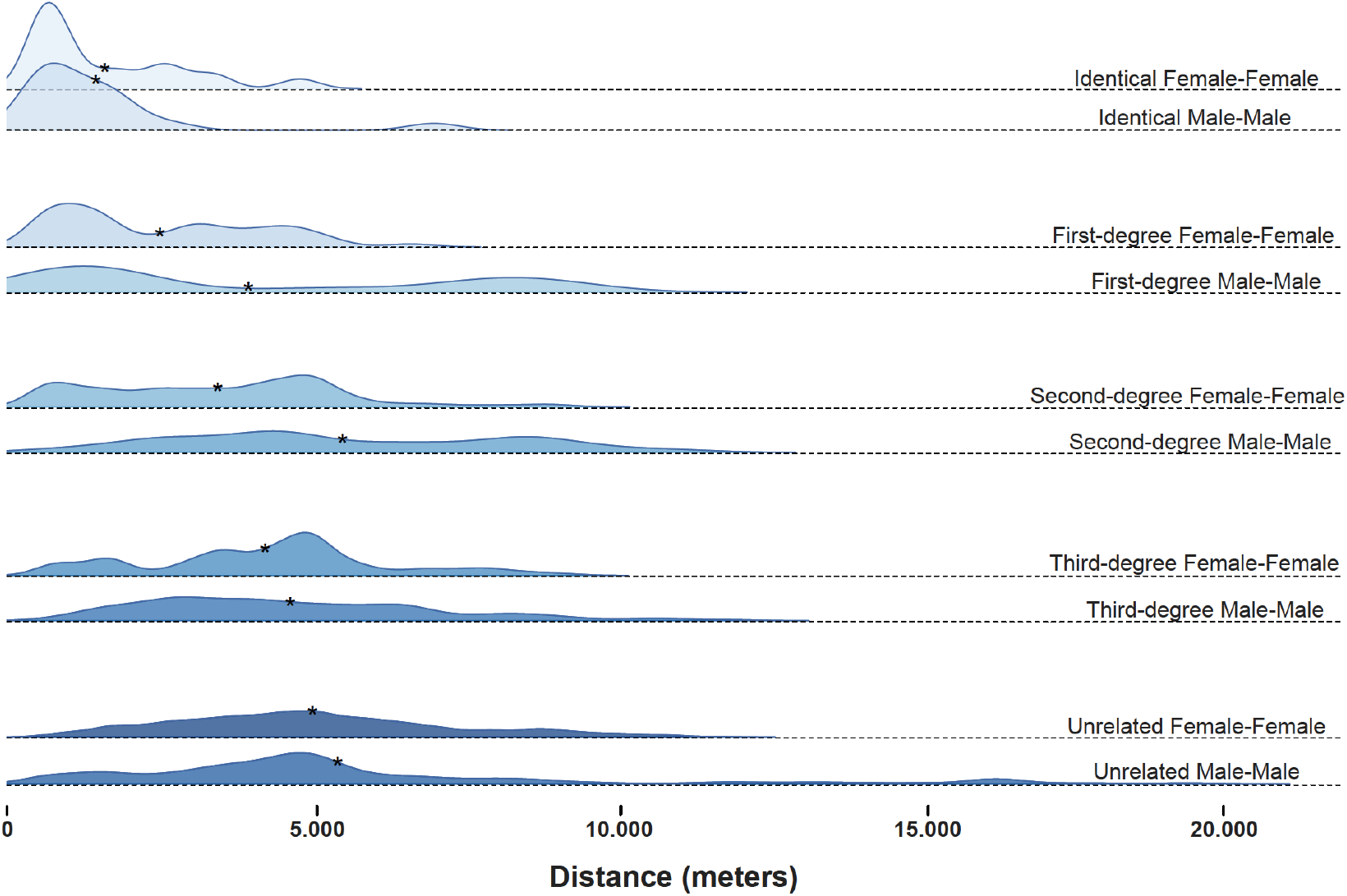
Distribution of Euclidean distances between geographic coordinates of samples, grouped by sex and degree of relatedness. Identical individuals are located significantly closer to each other compared to other categories. On average, related males are found farther apart than related females. Asterisks indicate the median of each distribution.

### 3.4. Population Size Estimation

By identifying identical samples through genetic matching, we determined the recapture rate of individuals. Of the successfully genotyped individuals, 55.5% were sampled once, 26.0% were sampled twice, and 18.5% were sampled three or more times (**Figure 8A**). Using these recapture frequencies, we simulated expected recapture distributions for various population sizes. The observed distribution most closely matched simulations for a population size of 164 to 208 individuals (95% CI) (**Figure 8B**).

**Figure 8.**
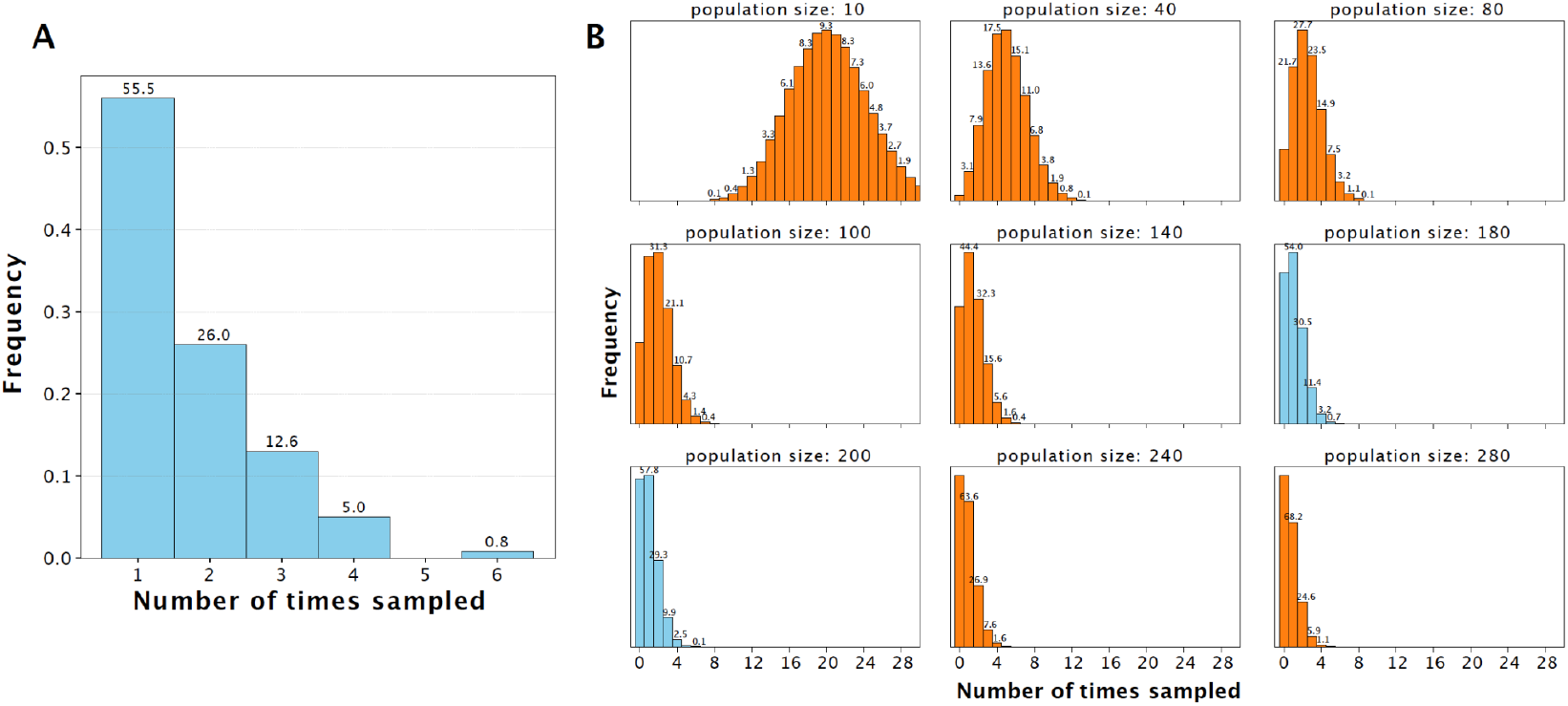
Non-invasive mark-recapture estimation of population size. **(A)** Frequency distribution of re-sampled individuals, showing that 55.5% of unique individuals were sampled once, 26.0% were sampled twice, and 18.5% were sampled three or more times. **(B)** Comparison of the observed re-sampling distribution to simulated distributions generated for different population sizes. The observed data most closely match simulations of a population size between 180 and 200 individuals.

### 3.5. Metagenomic Dietary Analysis

The detected plant DNA in the droppings confirms the reliance of adult capercaillie on key boreal forest species. The most abundant plant species in the samples was Vaccinium vitis-idaea (Lingonberry) and *Vaccinium myrtillus* (Blueberry), a known staple food source. DNA from *Alnus glutinosa* (Black Alder), *Picea abies* (Norway Spruce) and *Pinus sylvestris* (Scotch pine) was also identified, consistent with the consumption of buds and catkins, particularly during the spring (**Figure 9A**). Additionally, trace amounts of DNA from the lichen *Platismatia glauca* (Lungwort Lichen) were found, likely ingested incidentally along with other vegetation. The seasonal dynamics of plant consumption are illustrated in **Figure 9B**, which shows a heatmap of chloroplast DNA abundances across sampling months in 2022 and 2023. This analysis highlights temporal shifts in diet, with increased representation of coniferous species such as *Picea abies, Pinus sylvestris, and Larix sibirica* during spring, and higher abundances of berry plants such as *Vaccinium myrtillus* later in the season. We also recovered a broad spectrum of invertebrate DNA, highlighting the rich food sources available for capercaillies in Tyresta (**Figure 9A**). The findings include numerous species from orders known to be important for chick survival and growth. Beetles (Coleoptera): A diverse assemblage of beetle species was identified. This included various ground beetles (*Agonum fuliginosum, Harpalus rubripes, Pterostichus niger*), leaf beetles (*Agelastica alni, Chrysolina oricalcia*), and rove beetles (*Othius punctulatus, Philonthus spinipes*). The presence of both ground-dwelling and foliage-associated species could indicate that chicks forage in multiple microhabitats. Moths and Butterflies (Lepidoptera): The caterpillars of moths and butterflies are a primary food source for chicks. DNA from a wide variety of lepidopteran species was detected, including the Pine-tree Lappet (*Dendrolimus pini*), Nun Moth (*Lymantria monacha*), and Emperor Moth (*Saturnia pavonia*), whose caterpillars are large and protein-rich. Numerous other species, such as the Figure of Eight (*Diloba caeruleocephala*) and Buff-tip (*Phalera bucephala*), were also identified, demonstrating that chicks consume a broad range of available caterpillars. The analysis also revealed DNA from several other invertebrate groups. These included caddisflies (e.g., *Molanna angustata, Odontocerum albicorne*), mayflies (*Ecdyonurus torrentis*), dragonflies (*Sympetrum striolatum*), spiders (*Tetragnatha montana*), and earthworms (*Aporrectodea icterica, Lumbricus terrestris*). This diversity underscores the opportunistic foraging behavior of capercaillie, consuming a wide variety of small invertebrates as they become available in the forest ecosystem.

**Figure 9.**
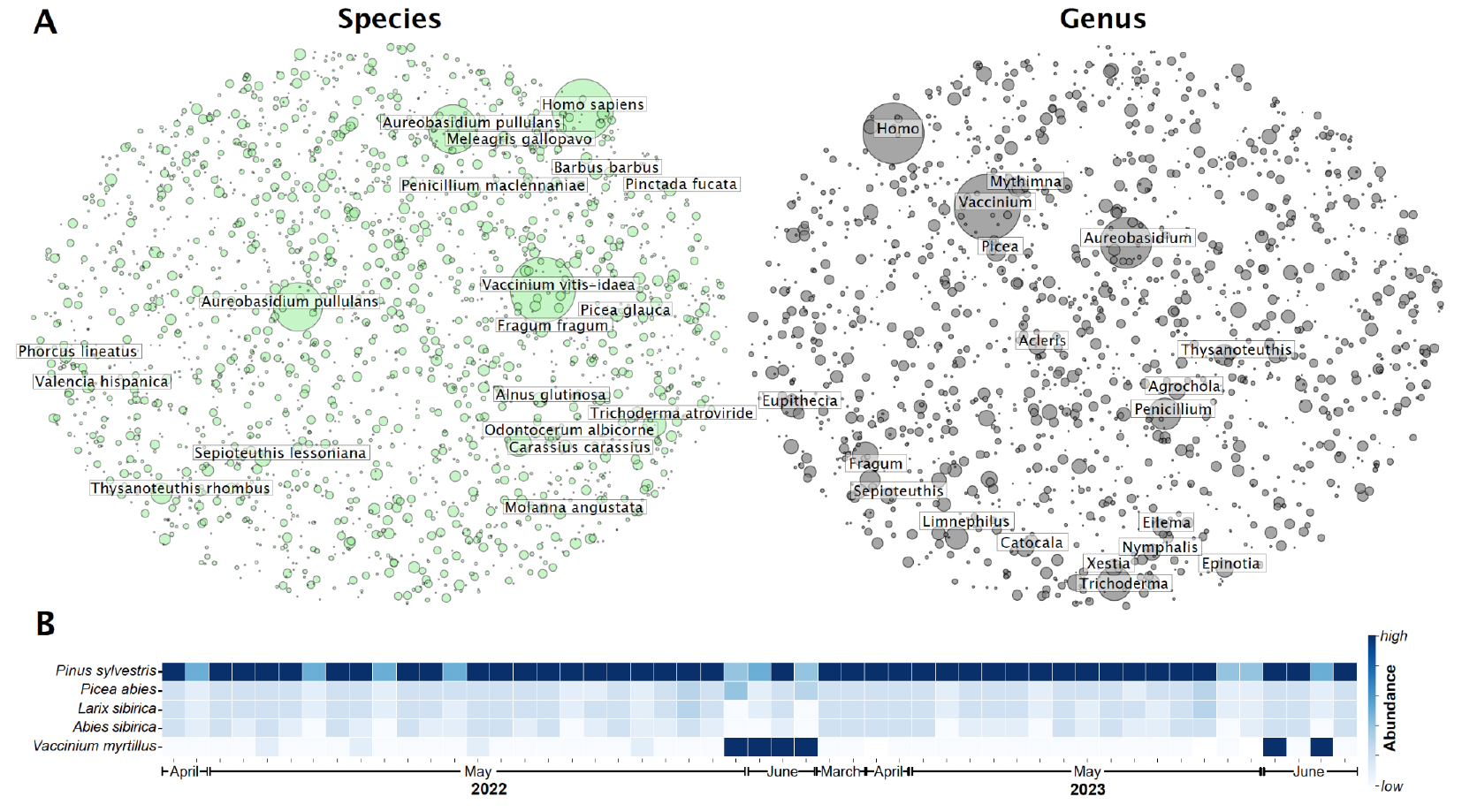
Metagenomic Profile of Capercaillie Diet and Gut Fauna. **(A)** These bubble plots visualize the taxonomic composition of non-host DNA found in capercaillie faecal samples, with the size of each bubble reflecting the relative abundance of reads from that taxon. The analysis is shown at the Species level (left) and Genus level (right). The plots reveal key dietary components such as *Pinus* (pine) and *Vaccinium* (Blueberry/cranberry), as well as various fungi and other microorganisms present in the birds’ environment or gut. Human DNA (*Homo sapiens*) was likely introduced during sample collection. **(B)** Seasonal variation in chloroplast abundance (coverage) for the most abundant plant species detected in the dataset across 2022 and 2023. Warmer colors indicate higher coverage and cooler colors indicate lower coverage.

## 4. Discussion

In this study we leverage the power of low-coverage whole-genome sequencing on non-invasive samples to conduct conservation assessment of the Western Capercaillie population in Tyresta National Park (Carroll et al. 2018). By combining genomic methods and metagenomic data, we estimated the population’s size, evaluated its genetic health and structure, reconstructed kinship patterns, and detailed its dietary habits (Kardos, Luikart, and Allendorf 2015). The findings reveal a population that, while currently maintaining a degree of genetic robustness, is small and isolated, facing vulnerabilities that require targeted conservation management (Frankham, Bradshaw, and Brook 2014; Keller 2002). Our results underscore the value of non-invasive genomic approaches for providing a holistic understanding of elusive wildlife populations.

### 4.1. The Power of Non-Invasive Genomics

A key methodological outcome of this research is the successful application of lcWGS to samples collected entirely non-invasively. We confirmed that shed feathers are a superior source of high-quality host DNA compared to faecal samples, consistently yielding higher coverage suitable for detailed genomic analyses like heterozygosity and ROH estimation. Faecal samples, while less effective for host genomics, proved to be a valuable source of metagenomic DNA, offering a window into the birds’ diet. This dual approach, using feathers for host genetics and faeces for ecological context, represents a powerful, cost-effective, and low-disturbance toolkit for the comprehensive monitoring of sensitive species like the capercaillie. The ability to reliably determine sex, identify individuals for mark-recapture estimates, and assess kinship without ever capturing a bird is a significant advantage for conservation practice.

### 4.2. A Small but Resilient Population on a Genetic Knife-Edge

Our genetic sampling simulations estimate the current population size at approximately 164–208 individuals. However, because our sampling protocol spanned two seasons, the assumption of a closed population is unlikely to hold. Demographic processes such as births, deaths, immigration, and emigration may have occurred during this period, potentially altering the proportion of re-sampled individuals. This could lead to an overestimation of the true population size. The reported estimate should therefore be regarded as an upper bound, with the actual population likely somewhat smaller. Translated into density, this corresponds to a maximum of 3.5–4.4 individuals per km² in Tyresta National Park. For a species with large territorial demands, this number is concerningly small, rendering the population highly susceptible to demographic stochasticity, environmental catastrophes (like fire or disease outbreaks), and the long-term erosion of genetic diversity. Despite this small size, the population’s current genetic health appears surprisingly sound. Genome-wide heterozygosity is comparable to that of other outgroup populations from mainland Europe, suggesting that the population has not yet passed through a severe, prolonged bottleneck that would have purged significant genetic variation. Similarly, the overall level of inbreeding, as measured by the total length of ROH, is relatively low. However, this optimistic picture is tempered by a warning sign: the presence of a few individuals with extremely long ROH segments. These segments are signatures of recent, close-kin mating (e.g., parent-offspring or full-sibling). While not yet systemic, such events can be a direct consequence of limited mate availability in a small, isolated population (Kardos, Luikart, and Allendorf 2015). An increase in the frequency of such matings could precipitate a rapid decline into inbreeding depression,leading to reduced fertility, compromised immune function, and lower survival rates. The Tyresta population is, therefore, balanced on a genetic knife-edge, currently healthy but with clear genomic indicators of future risk.

### 4.3. Population Structure: Isolation and Female-Biased Dispersal

The PCA and mtDNA analyses confirm that the Tyresta capercaillies function as a single, panmictic population, genetically distinct from a Swedish individual sampled in Jämtland and European populations. The finding that males, on average, have slightly more relatives within the park than females might suggest female-biased dispersal. This is a common pattern in avian species, where females typically disperse from their natal territories to find mates, while males remain more philopatric. Spatial analysis of relatedness further supports this view: most identical male samples were found within 1 km of each other, suggesting that this distance may approximate male territory size within the park. Nevertheless, a few identical males were found up to 5 km apart, indicating that some males either maintain unusually large ranges or occasionally disperse across lek sites. Field studies of capercaillie show clear differences in territory and home range size between males and females. Radio-tracking in Norway found that adult males typically occupy home ranges of a 0.5–1.5 km², occasionally exceeding 2.0–3.0 km², with much smaller defended display territories at leks measuring only a few tens of meters across (Per Wegge 1987; Rolstad and Wegge 1987). In central Europe, (Storch 1995) reported larger male ranges of 0.9–2.1 km² in Bavaria. Females generally use smaller areas, particularly during brood-rearing, with home ranges of 0.2–0.8 km² recorded in Norwegian studies (P. Wegge and Kastdalen 2007). Habitat selection work in Germany has shown that females’ ranges are closely tied to blueberry-rich forests and vary seasonally (Storch 1993). Overall, males occupy larger and more stable ranges centred on leks, while females shift habitats more dynamically, reflecting brood needs and resource availability.

More generally, our genetic data showed that related females tended to be located closer together than related males, consistent with the idea that male dispersal distances are larger within Tyresta. This dynamic has implications for the population’s management. Females are likely the primary conduits of gene flow. The few outlier individuals observed in the PCA are likely the result of successful immigration by females, an event that provides a vital infusion of novel genetic material. Therefore, conservation strategies must prioritize not only the habitat within the park but also the landscape matrix surrounding, including leks in the neighboring areas to sustain the entire population. Maintaining or creating functional habitat corridors that facilitate safe passage for dispersing females is arguably the most critical action for ensuring the long-term genetic viability of the Tyresta population (Whiteley et al. 2015).

The genetic data indicate a population density corresponding to a maximum of 3.5–4.4 individuals per km² in Tyresta National Park. This can be compared to estimates of Finnish late-summer (post-fledging) averages on the order of∼2–4 birds/km**²** (with wide spatial variation), while Norwegian spring densities commonly fall around∼1–3 birds/km² when sexes are combined, with site- and year-specific values ranging lower or higher depending on habitat and cycle phase (Odden et al. 2003; Finne et al. 2000; Miettinen et al. 2008; Brøseth, Kleven, and Bevanger 2025).

### 4.4. Linking Dietary DNA to Habitat

The identification of key plant species such as lingonberry, blueberry, Scots Pine, Norway Spruce and alder reaffirms the reliance of adult capercaillies on characteristic components of the boreal forest flora. The seasonal variation in detected chloroplast DNA further highlights how diet shifts across the year,with conifer buds and catkins (e.g., from Spruce and Pine) consumed predominantly in spring, and berry plants like *Vaccinium* dominating later in the season. Such patterns are consistent with long-standing ecological observations of capercaillie foraging behavior, but our results provide non-invasive direct molecular evidence of these dietary transitions. Equally important is the large diversity of invertebrates detected, spanning dozens of beetle, moth, caddisfly, mayfly, and other taxa. This broad spectrum underscores the availability of protein-rich food sources and the opportunistic foraging strategy of capercaillies. Importantly, capercaillie chicks are almost exclusively insectivorous during their first weeks of life, and their survival and growth are directly tied to the abundance of invertebrates, particularly caterpillars and beetles. Our findings therefore reinforce that capercaillie conservation cannot be separated from the conservation of invertebrate diversity. Taken together, these results demonstrate that a healthy capercaillie population depends on an ecologically complex forest ecosystem, one that provides a dynamic plant food base across seasons and sustains a rich invertebrate food web. Forest management practices must therefore aim to preserve this complexity.

### 4.5. Conservation Recommendations

In synthesis, the capercaillie population of Tyresta National Park is a small, isolated, and highly interconnected unit. It currently retains adequate genetic diversity but shows genomic warning signs of inbreeding. Its persistence is fundamentally linked to the integrity of its old-growth forest habitat and the rich biodiversity contained within it. The long term-effects of the large wildfire in 1999, which devastated the central part of the Park, are still evident on the capercaille population. It is therefore imperative that the habitats surrounding the national park, which are also sustaining a capercaille population, are considered in the long-term management of the capercaillie in the area and that in particular, leks and environments critical for chick survival are protected from intense forestry. We demonstrate that a non-invasive sampling effort, when analysed at genome-scale can provide a roadmap for the conservation of this population (Allendorf, Hohenlohe, and Luikart 2010; Shafer et al. 2015; van der Valk and Dalèn 2024).

## 5. Acknowledgments

We are grateful for the contribution of Bodil Ravn and Karin Asker Woodsack for collection of samples and Lina Ljungqvist, Magnus Marin and the Tyresta National Park rangers for their contribution to sample collection and logistics support for the project. Sequencing was performed by the SNP&SEQ Technology Platform in Uppsala. The facility is part of the National Genomics Infrastructure (NGI) Sweden and Science for Life Laboratory. The SNP&SEQ Platform is also supported by the Swedish Research Council and the Knut and Alice Wallenberg Foundation. The study was supported by a grant from Stiftelsen Alvins Fond (Naturvårdsverket) to Stiftelsen Tyrestaskogen.

## 6. Data availability

All sequencing data generated in this study is available at ENA under project ID: PRJEB88937

